## Supplementary figures and images for "Biological invasions alter environmental microbiomes: a meta-analysis"

### S1 Figure

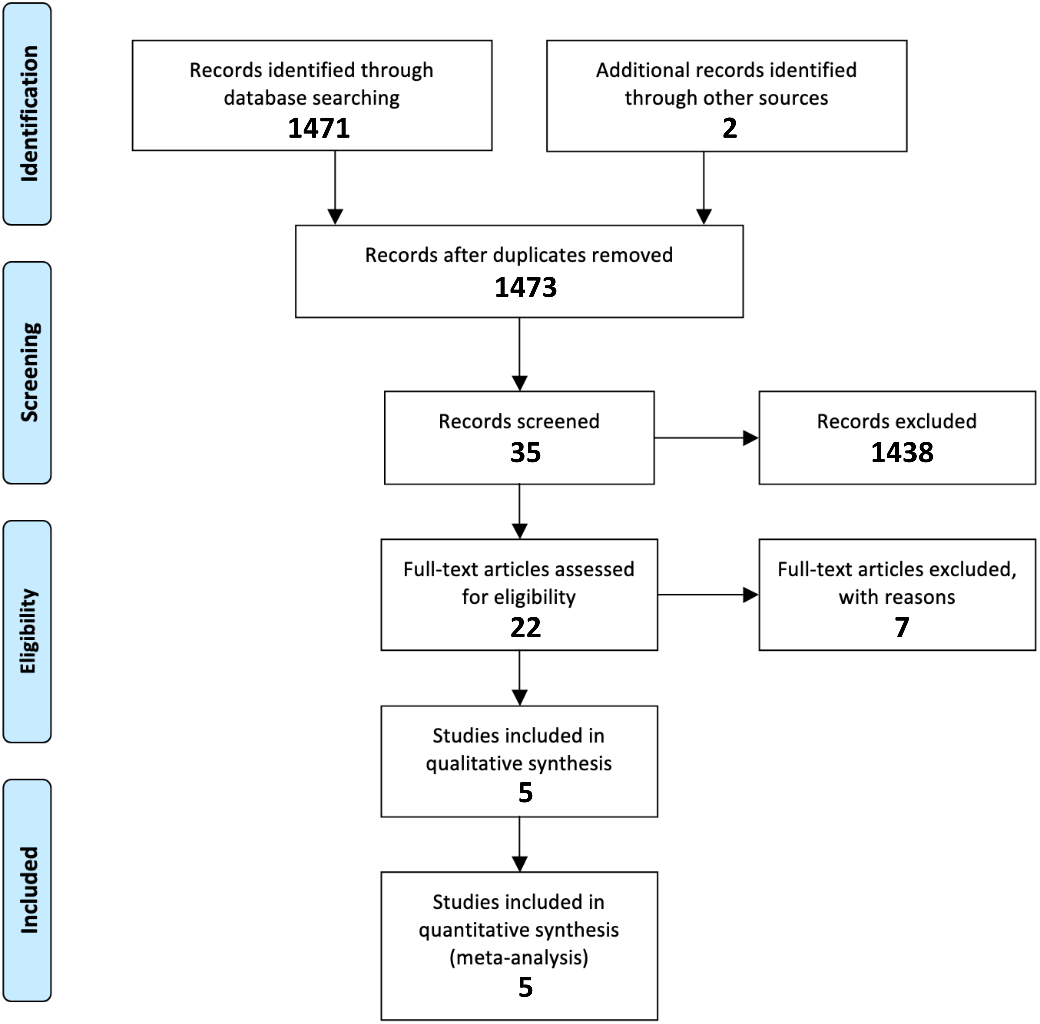
