## Supplementary material for "Biological invasions alter environmental microbiomes: a meta-analysis": S2 Table

**S2 Table.** Comparison of two different linear mixed-effects models testing the effect of biological *sample type* (invaded or control), *organism* (plant, mammal, mussel), and their interactions, on Shannon diversity index of the environmental microbiome. In Model 1 we included *studyID* as random factor, while in Model 2 we included both *studyID* and *environment* (soil or water) as random factors.

<table>
<tr><td colspan="4">Model 1, AIC = -160.88<br>lmer(Shannon ~ Sample_type * Organism_group * (1|Study_ID))</td></tr>
<tr><td></td><td>$\chi^2$</td><td>df</td><td>P</td></tr>
<tr><td>Sample_type</td><td>3.85</td><td>1</td><td>0.04</td></tr>
<tr><td>Organism_group</td><td>84.18</td><td>2</td><td><0.001</td></tr>
<tr><td>Sample_type* Organism_ group</td><td>3.51</td><td>2</td><td>0.17</td></tr>
<tr><td colspan="4"></td></tr>
<tr><td colspan="4">Model 2, AIC = -158.88<br>lmer(Shannon ~ Sample_type * Organism_group * (1|Study_ID) * (1|Environment))</td></tr>
<tr><td></td><td>$\chi^2$</td><td>df</td><td>P</td></tr>
<tr><td>Sample_type</td><td>3.85</td><td>1</td><td>0.04</td></tr>
<tr><td>Organism</td><td>59.7</td><td>2</td><td><0.001</td></tr>
<tr><td>Sample_type* Organism</td><td>3.51</td><td>2</td><td>0.17</td></tr>
</table>
