## Supplementary material for "Biological invasions alter environmental microbiomes: a meta-analysis": S3 Table

**S3 Table.** Comparison of two different PERMANOVA models testing the effect of biological *sample type* (invaded or control), *organism* (plant, mammal, mussel), and their interactions, on the structure of the environmental microbiome. In Model 1 we included *studyID* and *speciesID* as stratification factor, while in Model 2 we included *studyID*, *speciesID* and *environment* (soil or water) as stratification factors.

| **Model 1**<br>**~ Sample_type * Organism_group, strata = c("Study_ID", "Species_ID")** | | | |
|---|---|---|---|
| | **$R^2$** | **F** | **P** |
| *Sample_type* | 0.011 | 6.68 | <0.001 |
| *Organism_group* | 0.411 | 118.86 | <0.001 |
| *Sample_type* Organism_group* | 0.007 | 2.1 | 0.01 |
| **Model 2**<br>**~ Sample_type * Organism_group, strata= c("Study_ID", “Environment”, "Species_ID")** | | | |
| | **$R^2$** | **F** | **P** |
| *Sample_type* | 0.011 | 6.68 | <0.001 |
| *Organism_group* | 0.411 | 118.86 | <0.001 |
| *Sample_type* Organism_group* | 0.007 | 2.1 | 0.02 |
