## Supplementary material for "Biological invasions alter environmental microbiomes: a meta-analysis": S4 Table

**S4 Table.** Results from two different the linear mixed-effects model testing the abundance of each bacterial family against *sample type* (invaded or control), *organism* (plant, mammal, mussel), and their interactions. In Model 1 we included *studyID* as random factor, while in Model 2 we included both *studyID* and *environment* (soil or water) as random factors.

| | **Model 1*** | | **Model 2§** | |
|---|---|---|---|---|
| **Family** | **$\chi^2$** | **P** | **$\chi^2$** | **P** |
| Burkholderiaceae | 0.22 | 0.63 | 0.18 | 0.67 |
| Sporichthyaceae | 0.02 | 0.86 | 0.03 | 0.86 |
| Chthoniobacteraceae | **17.28** | **<0.001** | **17.28** | **<0.001** |
| Chitinophagaceae | **27.77** | **<0.001** | **27.78** | **<0.001** |
| Gemmataceae | **15.04** | **<0.001** | **17.42** | **<0.001** |
| Solibacteraceae (Subgroup 3) | **30.75** | **<0.001** | **30.68** | **<0.001** |
| Gemmatimonadaceae | **86.45** | **<0.001** | **86.46** | **<0.001** |
| Xanthobacteraceae | 1.23 | 0.26 | 1.22 | 0.27 |
| Ord. Tepidisphaerales | 3.80 | 0.05 | 3.76 | 0.05 |
| Sphingomonadaceae | **14.20** | **<0.001** | **14.21** | **<0.001** |
| Pyrinomonadaceae | 0.47 | 0.49 | 0.47 | 0.49 |
| Pedosphaeraceae | **12.09** | **<0.001** | **12.10** | **<0.001** |
| Solirubrobacteraceae | **10.36** | **<0.01** | **10.27** | **<0.001** |
| Acetobacteraceae | **33.30** | **<0.001** | **33.30** | **<0.001** |
| Ord. Solirubrobacterales | 2.74 | 0.09 | 2.71 | 0.10 |
| Blastocatellaceae | **32.42** | **<0.001** | **32.43** | **<0.001** |
| Pirellulaceae | **41.78** | **<0.001** | **41.10** | **<0.001** |
| Beijerinckiaceae | 4.01 | 0.05 | 4.01 | 0.05 |
| Haliangiaceae | 2.32 | 0.12 | 2.29 | 0.13 |
| Micromonosporaceae | **6.17** | **0.01** | **6.12** | **0.01** |
| Nitrosomonadaceae | **5.67** | **0.02** | **5.69** | **0.02** |

*lmer(Abundance ~ Sample_type * (1|Study_ID) * (1|Invasive_species))
§lmer(Abundance ~ Sample_type * (1|Study_ID) * (1|Environment) * (1|Invasive_species))
