## Supplementary material for "Biological invasions alter environmental microbiomes: a meta-analysis": S5 Table

**S5 Table.** Comparison of the relative proportion of each bacterial family between control and invaded environments. Differences are assessed using a linear mixed-effects model testing the normalized proportions of each bacterial family against *sample type* (Model 1 in S4 Table).

| Family | Control | | Invaded | | $\chi^2$ | P |
|---|---|---|---|---|---|---|
| | *Mean* | *SE* | *Mean* | *SE* | | |
| Acetobacteraceae | 0.0115 | 0.0004 | 0.0090 | 0.0004 | 33.3 | **<0.001** |
| Beijerinckiaceae | 0.0128 | 0.0002 | 0.0125 | 0.0003 | 4.01 | 0.05 |
| Blastocatellaceae | 0.0131 | 0.0005 | 0.0135 | 0.0005 | 32.42 | **<0.001** |
| Burkholderiaceae | 0.0315 | 0.0022 | 0.0404 | 0.0029 | 0.22 | 0.63 |
| Chitinophagaceae | 0.0499 | 0.0012 | 0.0527 | 0.0012 | 27.77 | **<0.001** |
| Chthoniobacteraceae | 0.0457 | 0.0016 | 0.0420 | 0.0015 | 17.28 | **<0.001** |
| Gemmataceae | 0.0290 | 0.0008 | 0.0253 | 0.0007 | 15.04 | **<0.001** |
| Gemmatimonadaceae | 0.0373 | 0.0013 | 0.0316 | 0.0011 | 86.45 | **<0.001** |
| Haliangiaceae | 0.0130 | 0.0003 | 0.0118 | 0.0004 | 2.32 | 0.12 |
| Micromonosporaceae | 0.0135 | 0.0004 | 0.0118 | 0.0004 | 6.17 | **0.01** |
| Nitrosomonadaceae | 0.0118 | 0.0002 | 0.0116 | 0.0003 | 5.67 | **0.02** |
| Ord. Solirubrobacterales | 0.0118 | 0.0004 | 0.0111 | 0.0004 | 2.74 | 0.09 |
| Ord. Tepidisphaerales | 0.0250 | 0.0008 | 0.0231 | 0.0010 | 3.8 | 0.05 |
| Pedosphaeraceae | 0.0186 | 0.0005 | 0.0179 | 0.0004 | 12.09 | **<0.001** |
| Pirellulaceae | 0.0122 | 0.0003 | 0.0132 | 0.0004 | 41.78 | **<0.001** |
| Pyrinomonadaceae | 0.0211 | 0.0008 | 0.0197 | 0.0008 | 0.47 | 0.49 |
| Solibacteraceae (Subgroup 3) | 0.0239 | 0.0009 | 0.0189 | 0.0007 | 30.75 | **<0.001** |
| Solirubrobacteraceae | 0.0113 | 0.0003 | 0.0103 | 0.0003 | 10.36 | **<0.01** |
| Sphingomonadaceae | 0.0255 | 0.0008 | 0.0267 | 0.0007 | 14.2 | **<0.001** |
| Sporichthyaceae | 0.0073 | 0.0019 | 0.0154 | 0.0027 | 0.02 | 0.86 |
| Xanthobacteraceae | 0.0275 | 0.0008 | 0.0238 | 0.0009 | 1.23 | 0.26 |
